# Individual differences in learning behaviours in humans: Asocial exploration tendency does not predict reliance on social learning

**DOI:** 10.1101/061473

**Authors:** Wataru Toyokawa, Yoshimatsu Saito, Tatsuya Kameda

## Abstract

A number of empirical studies have suggested that individual differences in asocial exploration tendencies in animals may be related to those in social information use. However, because the ‘exploration tendency’ in most previous studies has been measured without considering the exploration-exploitation trade-off, it is yet hard to conclude that the animal asocial ‘exploration-exploitation’ tendency may be tied to social information use. Here, we studied human learning behaviour in both asocial and social multi-armed bandit tasks. By fitting reinforcement learning models including asocial and/or social decision processes, we measured each individual’s (1) asocial exploration tendency and (2) social information use. We found consistent individual differences in the exploration tendency in the asocial tasks. We also found substantive heterogeneity in the adopted learning strategies in the social task: One-third of participants were most likely to have used the copy-when-uncertain strategy, while the remaining two-thirds were most likely to have relied only on asocial learning. However, we found no significant individual association between the exploration frequency in the asocial task and the use of the social learning strategy in the social task. Our results suggest that the social learning strategies may be independent from the asocial search strategies in humans.

## 1. Introduction

To find better behavioural options in foraging, mate choice, nest search, etc., group living animals can benefit from asocial information-gathering strategy (e.g., reinforcement learning rules (Sutton and Barto, 1998; Trimmer et al., 2012)) and from strategic use of social information (Boyd and Richerson, 1985; Laland, 2004). Although there has been much recent interest in the inter-individual variation in both asocial and social learning behaviour (Mesoudi et al., 2016; Reader, 2015), little is known about whether and (if so) how they associate with each other.

The adaptive reason of the individual differences in asocial exploration tendency might be the trade-off between exploration and exploitation. Given the limited time/energy budget, a single animal must strike the right balance between trying unfamiliar behaviours to sample information (i.e., ‘exploration’) versus choosing known best behaviour (i.e., ‘exploitation’) so as to improve the long-term net decision performance (Cohen et al., 2007; Hills et al., 2014). The optimal balance of exploration-exploitation depends on the costs and benefits of information gathering, which may differ between individuals. For example, an individual with poor information processing performance may have large costs of exploration, an individual with shorter expected life-span may benefit less from sampling more information, while an individual experiencing a temporary volatile environment may be forced to explore so as to update their knowledge (Reader, 2015).

On the other hand, the individual variation in reliance on social information might come from the balance of cost and benefit of copying others (Mesoudi et al., 2016). For instance, an individual possessing inaccurate private information will potentially incur a large cost if relying solely on the private knowledge and hence may tend to copy others more (e.g., ‘copy-when-uncertain’ (Laland, 2004; Rendell et al., 2011)), an individual living in a large group may benefit more from following the majority (King and Cowlishaw, 2007), while an individual faced with a highly volatile environment may rely more on private information due to the potentially large cost from copying an out-of-date behaviour (Aoki and Feldman, 2014).

The increasing body of empirical studies has suggested that the individual differences in the asocial exploration tendency might associate with those in the social information use (but see Webster and Laland (2015)). For instance, the individual exploration propensity negatively correlates with individual tendency of copying conspecifics in barnacle geese *Branta leucopsis* (Kurvers et al., 2010a, 2010b) and zebra finches *Taeniopygia guttata* (Rosa et al., 2012), while the opposite is true in three-spined sticklebacks *Gasterosteus aculeatus* (Nomakuchi et al., 2009) and great tits *Parus major* (Marchetti and Drent, 2000). The interesting question here would be why each of these species shows such individual correlations between asocial and social search behaviour. Several possible mechanisms may be conceivable to them. For example, environmental volatility may increase asocial exploration tendency while also decreasing copying tendency. On the other hand, a common cognitive ability underlying both asocial and social learning may generate a positive correlation between them (Mesoudi et al., 2016).

However, the term ‘exploration’ has been used rather loosely in the previous literature, and has been confounded with other personality traits (reviewed in Réale et al. (2007)), which might have contributed to somewhat incoherent previous findings on the relation between asocial exploration and social information use. Broadly speaking, more active, neophilic, or bolder individuals tend to be labelled as ‘explorative’ while more inactive, neophobic, or shyer individuals tend to be labelled as ‘unexplorative’ (Réale et al., 2007). However, it was untested whether more active individuals actually gather information more (i.e., explore more) during the learning process compared to inactive individuals. Also, from the viewpoint of the exploration-exploitation trade-off, more explorative (exploitative) individuals are not necessarily bolder (shyer): Especially in a changing environment, individuals who seldom explore (i.e., who exploit the same option for a long time) may also be seen as very ‘bold’ (e.g., Carere and Locurto, 2011; Groothuis and Carere, 2005; Koolhaas et al., 1999). Therefore, it is yet hard to conclude that animal asocial ‘exploration-exploitation’ tendency may be tied to the social information use. A more clear-cut measurement of exploration behaviour is needed.

In this study, we focused on human learning behaviour in a multi-armed bandit (MAB) problem, and saw whether the individual differences in asocial exploration tendency might predict the reliance on social learning. In the MAB task, individuals have multiple choice options, but at the outset they do not have exact knowledge of which option is the most profitable (Figure 1a). In every round, each individual has to make a decision whether to exploit (i.e., choosing the option that has higher estimated reward value as of that round; see Methods section) or to explore (i.e., choosing the other option with lower estimated reward value). Because the MAB problem embeds the exploration-exploitation trade-off in its heart (Sutton and Barto, 1998), it is a suitable test bed for unambiguously measuring exploration behaviour (Daw et al., 2006; Keasar et al., 2002; Racey et al., 2011; Toyokawa et al., 2014). Fitting a reinforcement learning model to each participant’s decision data (O’Doherty et al., 2003), we quantified each participant’s asocial exploration tendency.

In addition to the asocial situation where participants engaged in the MAB task alone (hereafter, ‘solitary task’), participants also played the MAB task in a pairwise situation (‘paired task’) in which they were able to observe the other participant’s choice (but not the peer’s earned payoff) displayed on the monitor. To examine whether the participants adopted social learning strategy in the paired task, we fitted several asocial-and social-learning models to each participant’s decision data, and then selected the most likely learning model individually. Also, we analysed each participant’s gaze movement measured by an eye-tracker in order to confirm the participant’s information use during the task. Finally, we examined whether the exploration tendency in the solitary task (i.e., asocial exploration) might predict the use of social learning strategy in the paired task.

## 2. Material and methods

### *2-1* *Participants*

Fifty-six right-handed undergraduate students were randomly selected from a subject pool at Hokkaido University in Japan to participate in the experiment. Of these 56 participants, 8 participants failed at eye tracker calibration, leaving us with 48 participants (24 females; Mean Age ± S.D. = 19.0 ± 0.90) to be included in data analysis. After the experimental session, participants received monetary rewards based on their performance in the experimental tasks as compensation for their participation (mean ± S.D. = 1253 ± 18.5 JPY).

### *2-2* *Task overview: the restless 2-armed bandit*

Participants performed a restless 2-armed bandit task on a computer screen (Figure 1a). We used a ‘2-armed’ task as the simplest case of a MAB. Each participant had to repeatedly choose between two slot machines. The participant’s goal was to maximise the total reward earned over a sequence of plays by deciding which machine to play for how many times.

Each round started with a 1-second interval during which a crossbar was shown at the centre of the grey background. After this interval, two boxes (i.e., slot machines) appeared on the left and right side of the screen. Participants chose the left box or right box by pressing the 'left' or 'right' key (respectively) on a keyboard with their right hand. Participants had a maximum of 2 seconds to make their choices (decision interval): If no choice was made during the decision interval, a 'TIME OUT' message appeared in the centre of the screen for 2.5 seconds to signal a missed round (average number of missed rounds per participant was 0.44 out of 80 rounds in the ‘first solitary task’, 0.52 out of 180 rounds in the ‘paired task’, and 0.15 out of 80 rounds in the ‘second solitary task’). If participants responded within 2 seconds, the frame of the chosen option turned to be bold for 1.5 + (2 − *response time*) seconds so as to confirm their choice, followed by a 1-second display of earned points in the chosen box. After showing the rewards (or showing ‘TIME OUT’), i.e., after 4.5 seconds from the outset of the decision interval, the next round started with a crossbar.

Each option yielded random points (50 points = 1 JPY) from a normal probability distribution unique to each box, rounded up to the next integer, or truncated to zero if it would have been a negative value (although this never happened). The mean values of the probabilistic payoff were different between the two options. Additionally, the mean values of payoff were changing during the task (Figure 1b). The standard deviations of the probabilistic payoff distributions were identical for both boxes and did not change during the task (S.D. = 10). Before each task started, participants were informed about the total number of rounds (80 rounds in the first and second solitary tasks, respectively; 180 rounds in the paired task) as well as about the possibility that the mean payoff from the options might change at some points during the task. However, they were not informed of the actual value of mean payoff from each slot, when or how it would actually change, or the exact rate at which the payoff points would be transformed into JPY after the experiment. To confirm that our results would not change in different bandits’ settings (e.g., mean payoff, pattern or schedule of payoff changing), we used four different settings (hereafter, conditions) for the paired task as a between-pair design, randomly assigned for each pair (Online Supporting Material Figure S2).

The computer-based task was constructed using Python with PsychoPy package (Peirce, 2007, 2009) with Tobii SDK 3.0 for Python. Python code used for the task is available from the corresponding author. Further details of the settings of each task are available in Online Supporting Material S1-1.

**Figure 1.**
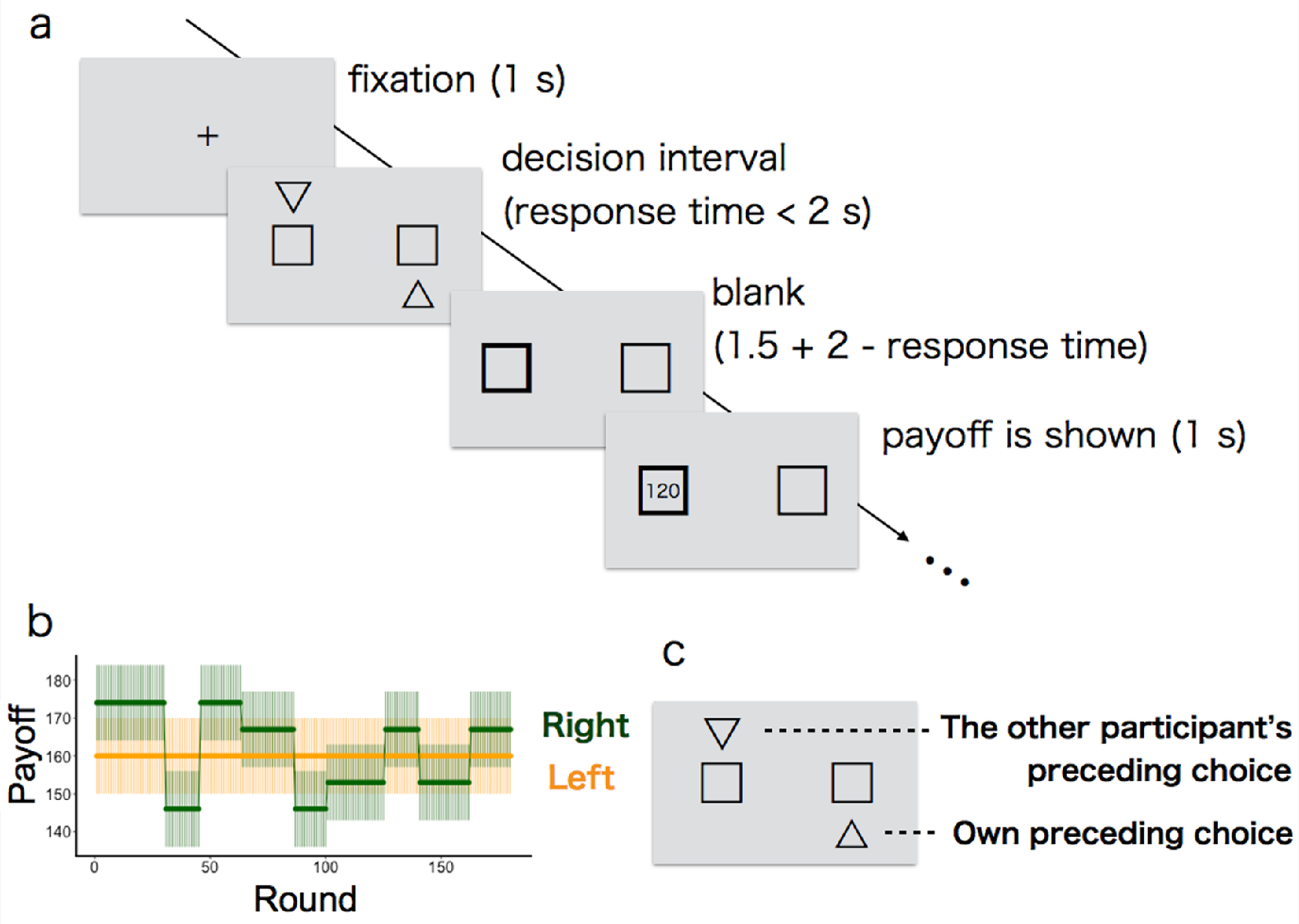
The restless 2-armed bandit task. (a) Illustration of the time line within a round. After fixation with a crossbar for one second, two slots (boxes) were presented. The participant had to choose one within 2 seconds (decision interval). When choosing one, the frame of the chosen option turned to be bold. Up to 3.5 seconds later, the number of payoff points earned was revealed to the participant (120 points in this example). After a further second, the next round started with a crossbar. (b) Example of mean payoffs for each option in paired task (bold lines). The payoff received for a particular choice is drawn from a Gaussian noise around each mean (shaded areas show 1 S.D. of this noise). Note that the most profitable slot (‘optimal option’) was switched several times due to the volatility; only one box was volatile and the other box’s mean was fixed. The left-right location of the volatile box was counterbalanced across pairs in the paired task (the right box was volatile in this example). (c) Example of social information in the paired task. The other participant’s choice in the preceding round was marked by a downward triangle (left in this example) while their own choice in the preceding round was marked by an upward triangle (right in this example).

### *2-3* *Experimental procedure*

For each experimental session, two participants of the same sex, randomly selected from the participant pool, were called to the laboratory. Upon arrival, each participant was seated in a separate soundproof chamber equipped with a computer terminal and received general instructions about the experiment. After filling in the consent form (see Ethics section), they read further paper-based instructions individually. Participants remained strictly anonymous to each other throughout the experiment.

A single experimental session was composed of three tasks: (1) first solitary task, (2) paired task, and (3) second solitary task. Although participants were informed that there would be three tasks in total, they did not know any details of each task until reading the instruction about the next task during the 5-minute break between the tasks. The three tasks differed in both the mean payoffs of slots and their changing pattern over time (see Online Supporting Material S1-1 for details). Furthermore, only in the paired task were participants able to see social information—the other participant’s choice made in the preceding round—in addition to their own preceding choice (Figure 1c). They were explicitly informed that both participants played the identical task so that they were able to understand that the social information could be informative. The session lasted for about 70 minutes in total.

### *2-4* *Acquisition and processing of gaze data*

We recorded each participant’s gaze movement during the task at 300 Hz using Tobii TX300 eye trackers connected to 23-inch monitors (Tobii Technology, Stockholm, Sweden). Participants were seated with their head positioned on a chinrest, 70 cm in front of the screen. Before each task, the calibration of the eye-tracker was validated with five fixation points and re-calibrated if needed.

We used Tobii’s default setting of noise reduction and fixation classification to process the gaze data. Because we were interested in participants’ information use before making each decision, we focused upon the gaze positions in the 1-second time window before each choice was made. Note that, because there was a 1-second interval between rounds, the 1-second time window did not include gaze positions in the previous round. The fixation positions of gazes were classified into the following five categories: *gazing-at-social-information, gazing-at-own-preceding-choice, gazing-at-the-left-box, gazing-at-the-right-box,* and *others* (see Online Supporting Material S1-2 for the gaze data processing). All eye-tracking data and Python code used in the gaze data processing are available from the corresponding author upon request.

### *2-5* *Testing exploration tendency*

In order to classify each choice in the solitary tasks as either exploration or exploitation, we fitted a standard reinforcement learning model to each participant’s choice data using maximal likelihood estimation method (Daw et al., 2006; O’Doherty et al., 2003). Following the previous empirical studies on social learning strategies in humans (McElreath et al., 2005, 2008), our learning model consists of two parts. First, the values for choosing the options (i.e., Q-values (Sutton and Barto, 1998)) are recursively updated by experienced reward according to the Rescorla-Wagner rule (Trimmer et al., 2012) that is commonly used in the animal learning literature. Second, the Q-values are transformed to the choice probability for each option by the ‘softmax’ choice policy (Daw et al., 2006). The mathematical expression of this asocial learning model is shown in the next sub-section (Eq.1 and Eq.2; see Online Supporting Material S1-3 for full details of the model fitting procedure).

After obtaining the most likely parameters of the asocial learning model separately for each participant, we classified each choice of each participant as either exploitative or explorative according to whether the chosen box had the larger Q-value (exploitation) or the smaller Q-value (exploration) between the two options. Then, we summed up the number of explorations for each participant for each of the solitary tasks. We calculated Pearson’s correlation coefficient between the numbers of explorations in the first and second solitary tasks so as to examine the individual consistency in exploration tendency.

### *2-6* *Determining learning strategies*

We examined four different learning models as candidates of participants’ strategies in the paired task: *asocial learning model* (AL), *unconditional-copying model* (UNC), *copy-when-uncertain model* (CWU), and *random choice model* (Random). AL and Random are asocial, while UNC and CWU are social learning models. Since each participant interacted with only one anonymous peer without seeing the peer’s earned payoff, we could not consider any ‘frequency based’ strategies (e.g., copy the majority) and ‘model based’ strategies (e.g., copy the prestigious individual or payoff-based copying) (Rendell et al., 2011). We fitted each of the models to each participant’s choice data individually.

All models have the same updating rule for Q-values, called the Rescorla-Wagner rule. Q-value for the left option is updated as follows:

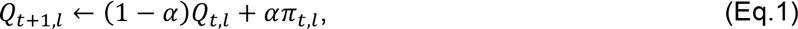

where *α* (0 ≤ *α* ≤ 1) is a parameter determining the weight given to new experience (learning rate) and *π_t,l_* is the amount of payoff obtained from choosing the left option (*l*) in round *t*. When the left option was not chosen in round *t*, the left option’s Q-value was not updated (*Q_t+1_,l* = *Q_t,l_*). The same updating rule applied to the right option. We set *Q*_0,*l*_ = *Q*_0,*r*_ = 0 because there was no reason to expect participants to have any prior preference for either option at the outset (McElreath et al., 2008).

Of course, we could have considered any other updating rules (e.g., Bayesian updating model (Payzan-LeNestour and Bossaerts, 2011)). However, because our scope here is examining whether the participants might conduct social learning or not, rather than quantifying detailed computational learning algorithms, we decided to focus only on the simple Rescorla-Wagner rule which has been shown to be evolutionary adaptive across a broad range of environmental conditions (Trimmer et al. 2012).

The Q-values were transformed into the choice probability in different ways between the four models, shown as follows.

#### *2-6-1* *Asocial learning model (AL)*

In the asocial learning model, the probability of choosing the left option in round *t* + 1 is given by the following softmax rule:

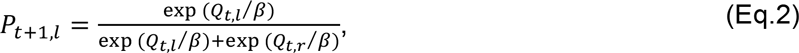

where *β* (*β* > 0) is a parameter that measures the influence of the difference between Q-values on choice (‘temperature’ parameter). As *β* → ∞, choice is completely random (i.e., *P*_*t*+1,*l*_ = 1/2; highly explorative). As *β* → 0, choice becomes deterministic, in favour of the option with the higher Q-value (i.e., highly exploitative). Therefore, *β* regulates the individual’s asocial exploration tendency.

This asocial learning model was also used in the last subsection (section 2-5) to estimate the exploration tendencies in the solitary tasks.

#### *2-6-2* *Unconditional-copying model (UNC)*

Next we considered unconditional/unselective copying, the simplest case of social information use (Laland, 2004). The individuals copied the other participant’s choice at the fixed rate *λ* (0 ≤ *λ* ≤ 1), otherwise their decisions were determined by the softmax rule (Eq. 2). In the case that the peer chose the left option in round *t*, the focal participant’s probability of choosing the left option at round *t* + 1 is given by:

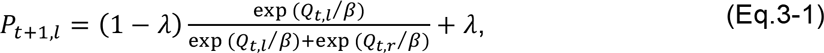

whereas when the peer chose the right option in round *t*, the participant’s probability of choosing the left option at round *t* + 1 is given by:

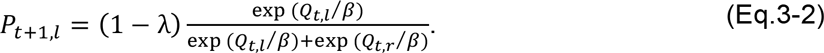

When *λ* = 1, the individuals always copy the peer’s choice. When *λ* = 0, the individuals rely only on asocial learning. We assumed that the individuals also relied only on the asocial learning rule when the peer missed the preceding round (i.e., when no choice was made by the peer).

#### *2-6-3* *Copy-when-uncertain model (CWU)*

A number of studies have suggested that animals are selective in timing to use social information, depending on the degree of uncertainty that they are experiencing (e.g., Coolen et al., 2003; Galef, 2009; van Bergen et al., 2004). To quantify the uncertainty level concerning which of the two slots is more rewarding at a given round, we used absolute difference in Q-values: The closer the Q-values between the two options, the higher the uncertainty. In the copy-when-uncertain model, when the peer chose the left option in round *t*, the focal participant’s choice probability for the left option at *t* + 1 is given by:

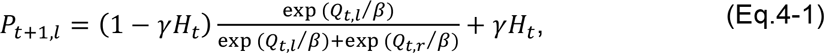

whereas when the peer chose the right option in round *t*,

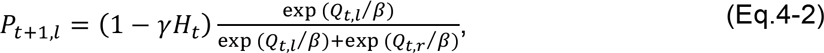

where

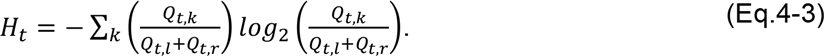

*γ* (0 ≤ *γ* ≤ 1) is a parameter that determines the upper limit of copying probability, and *H_t_* (i.e., information entropy, 0 ≤ *H_t_* ≤ 1) determines the actual copying rate at round *t*, where *k* ∈ {*l, r*}. When both Q-values are equal (*Q_l_* = *Q_r_*), the uncertainty becomes max (*H* = 1), leading the copying probability to be maximal (i.e., the individual copies the peer at the probability *γ* when uncertainty is the highest). As the difference between Q-values becomes larger, *H* approaches to 0, with the result that the choice is mostly determined by the asocial learning.

#### *2-6-4* *Random choice model*

We also considered the case that the choices are made randomly at fixed rate regardless of the Q-values, so as to verify that the participants did not behave just randomly in the experiment. The choice probability for the left option is always a fixed rate *ε* (0 ≤ *ε* ≤ 1); hence, the probability of choosing the right option is 1 − *ε*.

### *2-7* *Model selection*

We used Bayesian Model Selection (BMS) (Rigoux et al., 2014; Stephan et al., 2009) that estimates how likely it is that each learning model generated the data of a *randomly* chosen participant (see Online Supporting Material S1-4). Using BMS, we were also able to calculate the probability that each model had generated a *given* participant’s data (Stephan et al., 2009). Because here we were interested in interindividual variation of social information use, we mainly focused on the latter ‘individual-level’ probabilities for each model, rather than the former ‘group-level’ probability that BMS was originally aimed at. To verify this BMS estimation for each participant’s learning model, we also conducted the standard AIC comparison (see Online Supporting Material S1-5).

### *2-8* *The relation between gaze movement and learning strategy*

We examined the gaze movement so as to confirm the participants’ information use during the paired task. We focused on ‘*gazing-at-social-information before choice’* (yes = 1/no = 0) for each round as a binary response variable in a binomial generalised linear mixed model (GLMM), including the following three random effects: individuals, pairs, and bandit’s conditions. We considered estimated learning strategy (social = 1/asocial = 0), information uncertainty (i.e., similarity between Q-values [Eq. 4-3]), rounds, and possible 2-way interactions as fixed effects. Model selection was done based on each AIC value (Online Supporting Material S1-6).

### *2-9* *The relations between asocial exploration and social learning*

To examine whether the asocial exploration tendency might predict the use of social learning strategy, we analysed a binomial GLMM with social = 1/asocial = 0 learning as a binary response variable, with random effects of pairs and bandit’s conditions. We considered the asocial exploration tendency (frequency of explorative choices in the first solitary task), asocial learning performance (average payoff earned per choice in the first solitary task), and possible 2-way interactions as fixed effects. Model selection was done based on each AIC value (Online Supporting Material S1-7).

## 3. Results

### *3-1* *Individual consistency in the asocial exploration tendency*

The frequency of explorative choices in the first solitary task was positively correlated with that in the second solitary task (*r* = 0.58, *p* < 0.001; Figure 2). This result indicates that participants exhibited a stable asocial exploration tendency across the two solitary tasks. Results of the fitting of the asocial learning model are shown in Online Supporting Result S2-1.

**Figure 2.**
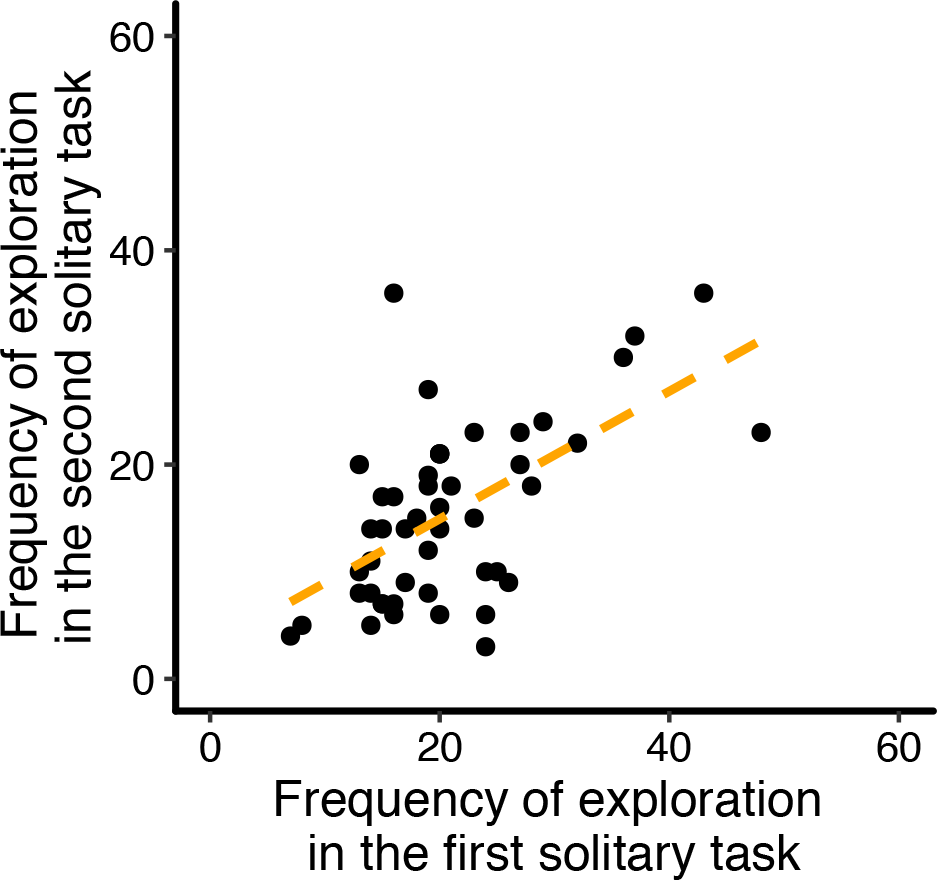
Individual consistency in the asocial exploration frequency. The x-axis shows the frequency of explorative choices in the first solitary task. The y-axis shows the frequency of explorative choice in the second solitary task. The dotted line is a linear regression.

### *3-2* *Detecting social learning strategies*

The Bayesian Model Selection (BMS) resulted in the heterogeneous distribution of learning strategies among the participants (Figure 3a). The estimated likelihood of each model for the whole data was 0.47 for the Asocial Learning model (AL), 0.10 for the Unconditional Copying model (UNC), 0.41 for the Copy-when-uncertain model (CWU), and 0.02 for the Random model.

Focusing on each individual's probabilities for each model, the BMS revealed that 32 participants were most likely to have adopted AL, while the remaining 16 participants were most likely to have adopted CWU (Figure 3b). No individuals were most likely to have used UNC or Random. We hereafter call the former 32 individuals ‘asocial learners’ and the latter 16 individuals 'social learners’ for simplicity. We also checked each model’s AIC values for each participant and confirmed that the result was not qualitatively changed (although three participants were most likely to have adopted the UNC model instead; see Online Supporting Material S2-2 and Figure S6).

Table 1 shows means (± 1 SDs) of best-fitted parameters for asocial and social learners. The mean value of *γ* (i.e., maximum copying probability) for social learners (CWU) was 0.162 (± 0.072), which means that, when uncertainty level was max (i.e., setting *Q*_*l*_ = *Q_r_* so that *H* = 1 and exp(*Q_t_*/*β*)/Σ exp(*Q_t_*/*β*) = 0.5, see Eq. 4), there was about 58% chance of choosing the same option as chosen by the other participant in the preceding round. Note that asocial learners should choose whichever option by 50% chance (i.e., randomly) when they have the maximum uncertainty. Therefore, there was at most 8% increase in the probability of “copying others’ choice” by social learners as compared to asocial learners. Together with the smaller frequency of social learners (N = 16) than asocial learners (N = 32), we will revisit this rarity of social learning in Discussion.

**Table 1.** Fitted parameter values (mean ± 1 S.D.) that provide the maximum likelihood for asocial learners (N = 32) and for social (copy-when-uncertain) learners (N = 16).

|  | *Asocial Learning*<br>*(AL)* | *Copy-when-uncertain*<br>*(CWU)* |
| --- | --- | --- |
| $\alpha$<br>(learning rate) | $0.842 \pm 0.187$ | $0.787 \pm 0.192$ |
| $\beta$<br>(temperature) | $11.31 \pm 10.39$ | $6.03 \pm 2.03$ |
| $\gamma$<br>(maximum copying<br>probability) | N.A. | $0.162 \pm 0.0720$ |
| N | 32 | 16 |

**Figure 3.**
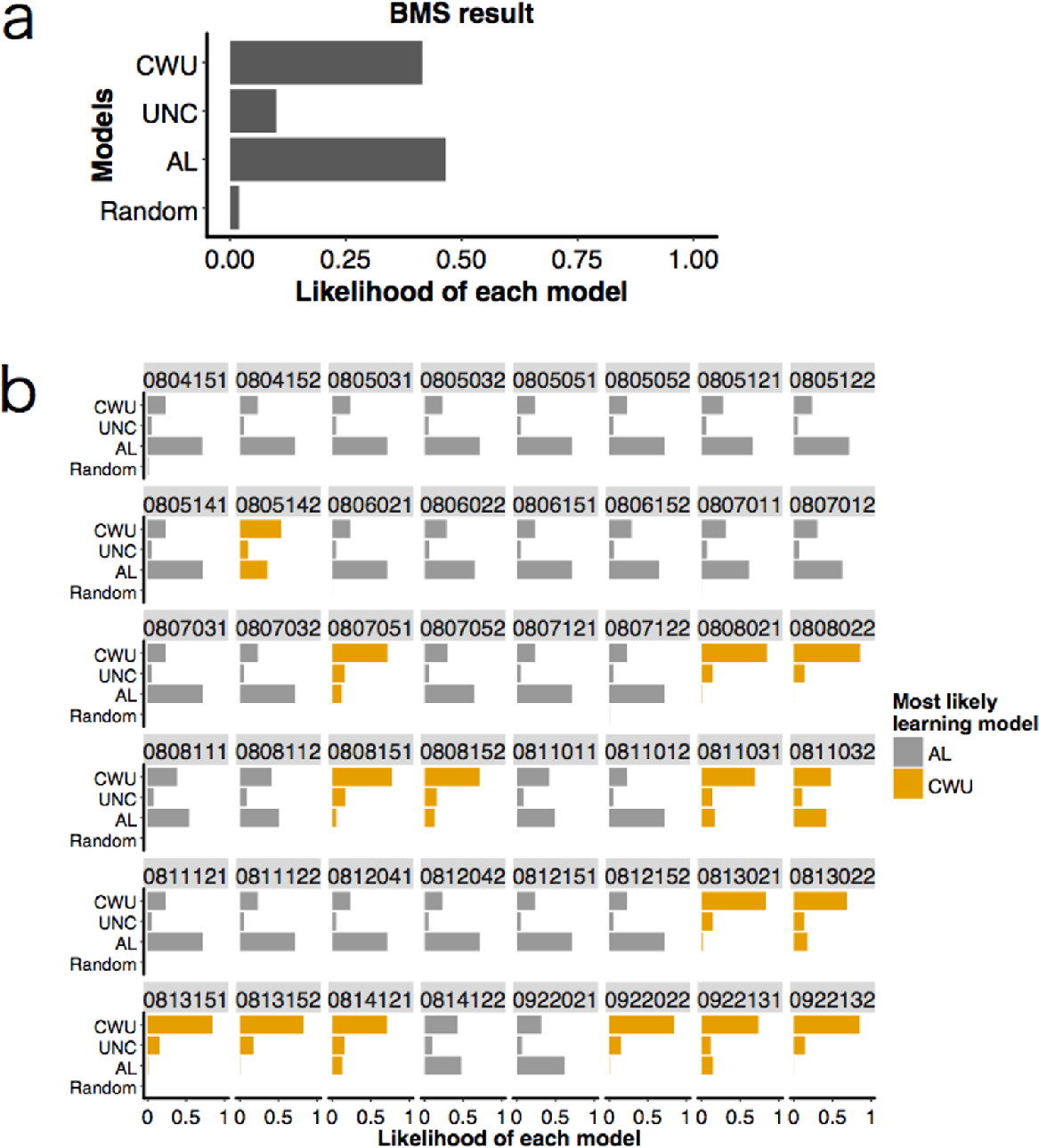
Results of Bayesian Model Selection. We considered the following four models: random choice model, asocial learning model (AL), unconditional copying model (UNC), and copy-when-uncertain model (CWU). (a) Distributions of probabilities concerning how likely it is that each model was adopted by a randomly chosen participant (i.e., group-level model likelihood). (b) Likelihood of each model for each participant (i.e., individual-level model likelihood). The numbers shown above each cell indicate participants’ IDs. Thirty-two participants were most likely to have used the asocial learning strategy (shown in grey); the remaining 16 participants were most likely to have used the copy-when-uncertain strategy (shown in orange).

### *3-3* *Social learning did not improve the performance*

Given the result that some participants used social learning, an interesting question is whether social learners performed better than asocial learners in the paired task. We compared each individual’s average payoff earned per choice using a Gaussian-GLMM with a fixed effect of learning type (asocial = 0/social = 1), and with two random effects of pairs and bandit’s conditions. The fixed effect was not significant, implying that social learners performed no better than asocial learners (*coefficient* [95% confidence interval] = 0.38 [−0.33, 1.12], *p* = 0.28; see Online Supporting Figure S8).

### *3-4* *The relation between gaze movement and learning strategy*

The result of BMS could be argued as an artefact of the limited number of models considered. To confirm that estimated ‘social learners’ actually saw the social information, we also examined the participants’ gaze patterns. We used the estimated learning strategy as a dummy fixed effect (i.e., asocial learner = 0; social learner = 1). The binomial-GLMM for the probability of looking at social information revealed the significant effects of uncertainty (coefficient [odds ratio] = −0.63 [0.53], *p* = 0.036), round (coefficient [odds ratio] = 0.24 [1.25], *p* < 0.001), the interaction between uncertainty and learning type (coefficient [odds ratio] = 1.45 [4.25], *p* = 0.0028), and the intercept (coefficient [odds ratio] = −3.25 [0.039], *p* < 0.001). However, the fixed effect of learning strategy itself was not significant (coefficient [odds ratio] = −0.65 [0.52], *p* = 0.37). The confidence intervals are shown in Online Supporting Table S2.

The significant negative effect of uncertainty suggests that the participants tended not to look at the social information when uncertainty (here defined as the closeness of the two options in terms of Q-values) was high. However, the significant positive interaction between uncertainty and learning strategies suggests that, when uncertainty was high, the social learners tended to look at the social information more frequently than asocial learners. This interaction effect is thus consistent with the behavioural-choice pattern predicted by the CWU model as compared to the AL model.

### *3-5* *The relation between asocial exploration tendency and social learning strategy*

Our results showed that (1) there were consistent individual differences in asocial exploration tendency across the two solitary tasks and that (2) use of social learning was heterogeneous between individuals. Given this, we finally examined whether the asocial exploration tendency in the solitary task might predict the use of social learning strategy in the paired task. We analysed a binomial GLMM that predicts the use of social learning strategy in the paired task (Online Supporting Material S1-7).

The selected GLMM contains both fixed effects of exploration frequency at the first solitary task and performance at the first solitary task. However, none of them were statistically significant (exploration tendency: coefficient [odds ratio] = −1.21 [0.30], *p* = 0.16, performance: coefficient [odds ratio] = −1.05 [0.35], *p* = 0.07). Hence, we cannot say either of them predicts a participant’s reliance on social learning. Looking at the scatter plots (Online Supporting Material Figure S8) rather than just considering p-values of the GLMM parameters, however, the average performance of the social learners in the first solitary task was not higher than the expected performance from completely random choices (Student’s t-test: *t_15_* = 0.64, *p* = 0.53), while that of asocial learners was better than the chance-level (Students’ t-test: *t_31_* = 6.15, *p* < 0.001; Figure S8a). It might suggest that the asocial learning performance in the first solitary task has a negative effect on the use of social learning in the paired task. On the other hand, the frequencies of explorations in the first solitary task were not different between asocial and social learners (Figure S9).

## 4. Discussion

In this study, we investigated human search strategies in the asocial/social 2-armed bandit tasks, respectively, and tested whether the individual differences in asocial exploration tendency in isolated settings might predict the use of social learning in group settings.

Across the first and second solitary tasks, our results showed the consistent individual differences in asocial exploration tendency (Figure 2). Since participants were not informed how the environmental change would occur in advance, it was virtually impossible for them to calculate the optimal exploration schedule (Gittins et al., 2011). Therefore, individual differences in calculation abilities are unlikely to explain this individual variation. Instead, it is known that human exploration tendency in a learning/decision task may be dependent on dopaminergic functions in the prefrontal cortical region of the brain (Daw et al., 2006; Frank et al., 2009), which is associated with tracking informational uncertainty (Yoshida and Ishii, 2006). Sensing decision uncertainty accurately is important in deciding when to explore in multiarmed bandit problems (Cohen et al., 2007). The individual differences in asocial exploration tendency shown in our result may be related to such individual differences in neural activity for sensing uncertainty, although we did not investigate direct neural mechanisms here.

As for the social task, we found a heterogeneous distribution of learning strategies: Two-thirds of the participants seems to have used the *asocial learning* strategy while the rest seems to have used the *copy-when-uncertain* strategy (Figure 3b), even though a few of them might have adopted the *unconditional-copying* strategy instead (Online Supporting Figure S6). The copy-when-uncertain type of social information use is predicted by adaptive evolutionary models (e.g., Boyd and Richerson, 1988), and has been repeatedly reported in empirical studies about non-human animals (e.g., Galef et al., 2008; Kendal et al., 2009) as well as humans (e.g., Kameda and Nakanishi, 2002; Morgan et al., 2011; Muthukrishuna et al., 2015). Our result replicated those findings. However, in the previous empirical studies, uncertainties were externally manipulated by, for example, varying the task difficulty (Morgan et al., 2011), changing the option numbers (Muthukrishuna et al., 2015), or changing costs required for getting asocial information (Kameda and Nakanishi, 2002). Instead of fixing such parameters via experimental manipulations, our copy-when-uncertain model tracks each individual’s internal information uncertainty (i.e., entropy) that can dynamically change during the course of learning. Our result thus provides a finer picture about possible plasticity in human learning strategies, shedding light on the individual flexibility of social information use during the course of problem solving.

It is noteworthy that, however, there were other forms of uncertainty that we did not consider here. For example, there were uncertainties concerning how noisily the payoff would be generated from a slot machine (i.e., variance of the payoff distribution), how often the environment would change (i.e., ‘unexpected uncertainty’), and what kind of distribution payoff was generated from (i.e., ‘structural uncertainty’ or ambiguity) (Cohen et al., 2007; Payzan-LeNestour and Bossaerts, 2011; Payzan-LeNestour et al., 2013). Further study is needed to investigate how those different types of uncertainty may affect the timing of social information use.

Technically, we used AIC as an approximation to the log-evidence for the Bayesian Model Selection (see Online Supporting Method). Compared to other forms of calculation (e.g., Free Energy approximation), AIC tends to prefer complex models because of the weak penalty for having more parameters (Penny, 2012; Stephan et al., 2009). It might thus have caused over-estimation of the frequency of social learners because our social learning models have more parameters than do asocial models. Nevertheless, the gaze movement patterns were consistent with the behavioural-choice pattern from the BMS result—individuals categorised as copy-when-uncertain learners saw the social information more often than asocial learning individuals when they faced high uncertainty (Result 3-4). Importantly, the BMS only considered the behavioural (i.e., choice) data, which was measured independently from the eye-tracking. Therefore, the result confirmed that we successfully captured the significant pattern of individual differences in learning strategies.

Although evolutionary theory generally tends to suggest heavy reliance on social learning in a broader range of situations (Rendell et al., 2011), only one-third of participants seemed to have used social information in our experiment. In addition to the low prevalence of social learners among participants, the maximum copying probability of social learners was also low (at most 8% higher than that of asocial learners). This rarity of social learning might be because social learning was not so useful in the paired task (Result 3-3). Social information is useful if and only if others’ behaviours can filter better options (Rendell et al., 2010). In our paired task, however, there was only one other individual playing the same task. Thus, the filtering effect from social learning was minimal. Also, there might be no ‘worth-copying’ peer (e.g., expert or veteran) because both participants started the task at the same time: When a focal participant was naïve to the task, so was the other participant. Therefore, information about the peer’s choice might not be so accurate compared to the own (asocial) learning experiences. Additionally, a recent empirical study suggests that the reliance on social information becomes stronger as the number of choice options increases (Muthukrishuna et al., 2015). Having only two options, therefore, our current study might underestimate the potential use of social learning. Further studies are needed to investigate whether social information use may change with group size and/or number of options.

We also explored the possible associations between behaviour in the solitary task and social learning in the paired task. Different from the previous empirical findings about non-human animals, we found no relation between asocial exploration tendency and the reliance on social learning. One possible reason for this difference might come from the inconsistent definition of ‘exploration’ in the previous literature. As described in Introduction, asocial exploration tendency has often been confounded with other personality traits that could relate to more general asocial learning ability (Réale et al., 2007). Better asocial learners may show less social information use because they possess more accurate private information (e.g., Kurvers et al., 2010a, 2010b), while the opposite (better asocial learners rely more on social learning) could also be plausible if both asocial and social learning reflect a common basic cognitive ability (Mesoudi et al., 2016). Indeed, a number of studies have shown that asocial learning ability correlates with the use of social information (Bouchard et al., 2007; Katsnelson et al., 2011; Mesoudi, 2011). Although it was not statistically significant, our result might also suggest that better asocial learners tend to ignore social information (Result 3-5; Figure S8a). Importantly, in our task, the asocial exploration tendencies were not correlated with the asocial learning performance (first solitary task: *r* = −0.20, *p* = 0.17; second solitary task: *r* = −0.14, *p* = 0.35; Online Supporting Figure S7). Our results that the asocial learning performance might have a negative effect on the use of social learning (but not exploration tendency) may suggest the importance of drawing a distinction between information-gathering behaviour (e.g., exploration tendency) and learning performance.

Overall, our study supports that humans are very selective about when to use social information. On the other hand, we should also acknowledge that our experimental set-up did not allow considering any ‘copy-from-whom’ (Heyes, 2015; Laland, 2004; Rendell et al., 2011) types of strategies, which might cause overlooking the potential social information use. Comparing ‘who’ strategies with ‘when’ strategies in the same framework will provide more comprehensive understanding on social learning for both human and non-human animals. We also believe that the computational learning model can be a strong tool for quantitative empirical investigations on animal social learning strategies (Daw et al., 2006; McElreath et al., 2005, 2008; O’Doherty et al., 2003; Sutton and Barto, 1998).

## 5. Ethics

This study was approved by the Institutional Review Board of the Centre for Experimental Research in Social Science at Hokkaido University (No. H26-01). Written informed consent was obtained from all participants before beginning the task.

## 6. Data accessibility

Behavioural data are available in Online Supporting Materials

## 8. Authors’ contributions

WT designed the study, performed the experiment, carried out the data analysis and drafted the manuscript; YS carried out the acquisition of gaze data and participated in the data analysis; TK participated in the design of the study, supervised the study and participated in writing the manuscript. All authors gave final approval for publication.

## 9. Acknowledgments

We are grateful to Shinsuke Suzuki for valuable discussions related to the computational learning models, and Mike Webster for helpful comments on earlier draft of this manuscript.

